# Novel viral diversity and geographic structuring in the New Zealand West Coast blackfly (*Austrosimulium ungulatum*)

**DOI:** 10.64898/2026.11.08.765125

**Authors:** Kimiya Byrne, Jessica A. Darnley, Stephanie J. Waller, Jemma L. Geoghegan

## Abstract

**Background:** Vector-borne diseases represent a major global public health burden, with climate change, land-use change and increasing human activity altering arthropod vector distributions and creating opportunities for infectious disease emergence. In Aotearoa New Zealand, insects commonly referred to as “sandflies” are blackflies belonging to the family Simuliidae. The West Coast blackfly, *Austrosimulium ungulatum*, is an endemic, human-biting species abundant throughout the South Island, yet its associated viral diversity remains largely unexplored.

**Methods:** Here, we used metatranscriptomic sequencing to characterise the virome of the West Coast blackfly and investigate patterns of viral diversity and community composition across geographical and environmental gradients. A total of 425 blackflies were collected from 19 locations spanning approximately 187 km.

**Results:** We identified 94 viral sequences across 16 orders and six phyla, all of which were putatively novel viruses. These included divergent lineages within families containing recognised vertebrate and arthropod pathogens, such as the *Rhabdoviridae* and *Orthomyxoviridae*. Viral community composition varied among sampling locations, but no significant associations were detected between land-use categories and viral diversity or community composition. In contrast, we revealed a weak but significant positive correlation between geographical distance and viral community dissimilarity, suggesting some degree of spatial structuring across the sampling range.

**Conclusions:** This study provides the first metatranscriptomic characterisation of the virome associated with the endemic West Coast blackfly, revealing substantial previously undescribed viral diversity and providing insights into the geographical distribution of arthropod-associated viral communities.

## Introduction

Vector-borne diseases pose a growing threat to human and animal health, driven by environmental change, expanding vector distributions and increasing global connectivity (1–3). Climate change is altering the geographical ranges, abundance and seasonal activity of arthropod vectors, while land-use change, urbanisation and agricultural expansion are reshaping interactions among vectors, wildlife, livestock and humans (4, 5). These changes have intensified interest in the diversity, ecology and evolutionary origins of arthropod-associated viruses, particularly those with the potential to cross host boundaries. Yet, despite major advances in virus discovery, our understanding of the RNA virosphere remains heavily biased towards a relatively small number of medically and economically important arthropod species (6). The vast majority of blood-feeding insects, including many that are abundant, geographically widespread or endemic to distinct ecosystems, remain largely unexplored.

Arthropods harbour an extraordinary diversity of RNA viruses and have played a fundamental role in their long-term evolution (7). Metatranscriptomic studies have revealed thousands of previously undescribed viruses across diverse invertebrate taxa, substantially expanding the known evolutionary breadth of the RNA virosphere and identifying deep evolutionary relationships between invertebrate- and vertebrate-associated viruses (7–9). These discoveries suggest that arthropods represent an important component of global viral biodiversity, extending well beyond their established roles as vectors of disease. However, the ecological processes governing the composition and distribution of arthropod viromes remain poorly resolved. In particular, the relative contributions of host identity, environmental conditions and geographical separation to viral community structure are incompletely understood. Furthermore, the recovery of viral sequences from whole insects does not necessarily indicate infection, as viruses may originate from blood meals, associated microbiota, parasites or other environmental sources (8, 10). Distinguishing these associations is essential for understanding the ecological and evolutionary significance of newly discovered viruses.

Research on blood-feeding arthropods has overwhelmingly focused on mosquitoes, despite the ecological diversity and potential epidemiological importance of other dipteran groups, including blackflies, biting midges and phlebotomine sandflies (11, 12). These insects occupy diverse habitats, often occur at high densities and interact with multiple vertebrate host species, creating opportunities for exposure to viruses circulating across different ecological communities (13, 14). Blackflies (Simuliidae), in particular, are globally distributed blood-feeding insects with complex aquatic and terrestrial life histories. Several species are established vectors of parasites, including Onchocerca volvulus, the causative agent of human river blindness, and avian Leucocytozoon species (15–19). Blackflies have also been associated with viruses such as eastern equine encephalitis virus, Venezuelan equine encephalitis virus, Rift Valley fever virus and vesicular stomatitis viruses, although evidence of their capacity to transmit these viruses remains limited or varies among species (19–21). Despite these associations, the broader RNA viral diversity of blackflies remains poorly characterised, particularly outside regions where they are recognised as vectors of human or animal pathogens.

Aotearoa New Zealand provides a distinctive setting in which to investigate arthropod-associated viral diversity. Its geographical isolation and high levels of endemism have shaped unique insect communities, many of which remain largely unexplored from a virological perspective (22). The insects commonly known as sandflies or namu in New Zealand are blackflies belonging to the endemic genus *Austrosimulium* (Simuliidae), rather than the phlebotomine sandflies (Psychodidae) that transmit pathogens such as Leishmania parasites and Toscana and Chandipura viruses elsewhere in the world (23–27). The genus *Austrosimulium* comprises 19 recognised species, three of which bite humans (27). Among these, the West Coast blackfly (*Austrosimulium ungulatum*) is particularly abundant across the west coast of the South Island, where it is a widespread biting nuisance. These insects occupy environments characterised by substantial variation in climate, topography and habitat, and feed on vertebrate hosts including birds and marine mammals. Despite their abundance and frequent contact with humans and wildlife, New Zealand blackflies are not known to transmit pathogens of public health importance (28).

The viral diversity associated with *Austrosimulium* remains almost entirely unknown. The only previous virological investigation of New Zealand blackflies used DNA sequencing to identify single-stranded DNA viruses, leaving their RNA viromes unexplored (28). This represents a substantial gap in our understanding of viral diversity in an endemic, blood-feeding insect lineage. More broadly, investigating viruses in geographically isolated arthropod populations offers an opportunity to explore how viral communities are assembled and maintained across heterogeneous environments. Whether viral diversity in these insects is structured primarily by local environmental conditions, geographical separation or broader ecological processes remains unknown.

Here, we used metatranscriptomic sequencing to characterise the RNA virome of the West Coast blackfly across 19 locations spanning the West Coast of the South Island, Aotearoa New Zealand. We investigated the diversity and evolutionary relationships of associated viruses and examined how viral community composition varied across geographical and environmental gradients. By exploring the virome of an endemic insect lineage that has received little virological attention, this study provides new insights into the hidden diversity of arthropod-associated RNA viruses and the ecological factors that shape their distribution.

## Methods and materials

### Ethics and sampling permissions

Under New Zealand legislation, the collection and use of most invertebrates for research do not require animal ethics approval (29). Accordingly, ethical approval was not required for the collection of blackflies (*Austrosimulium* spp.) in this study.

### Sample collection

Blackflies were collected from 20 sampling sites across 19 locations along approximately 187 km of the West Coast of the South Island, Aotearoa New Zealand, extending from the Blue Pools near Makarora to Hokitika Gorge (Figure 1). Sampling encompassed a range of environments, including coastal habitats, native forest and riverine ecosystems. A total of 425 blackflies were collected in November 2025 under relatively mild, humid and low-wind conditions favourable for blackfly activity. Specimens were collected using Biogents BG-Pro traps (Biogents, Germany), which use an attractant and fan-assisted collection system, or manually using sweep nets and forceps in areas of high blackfly abundance. At least four individuals were collected per site.

**Figure 1.**
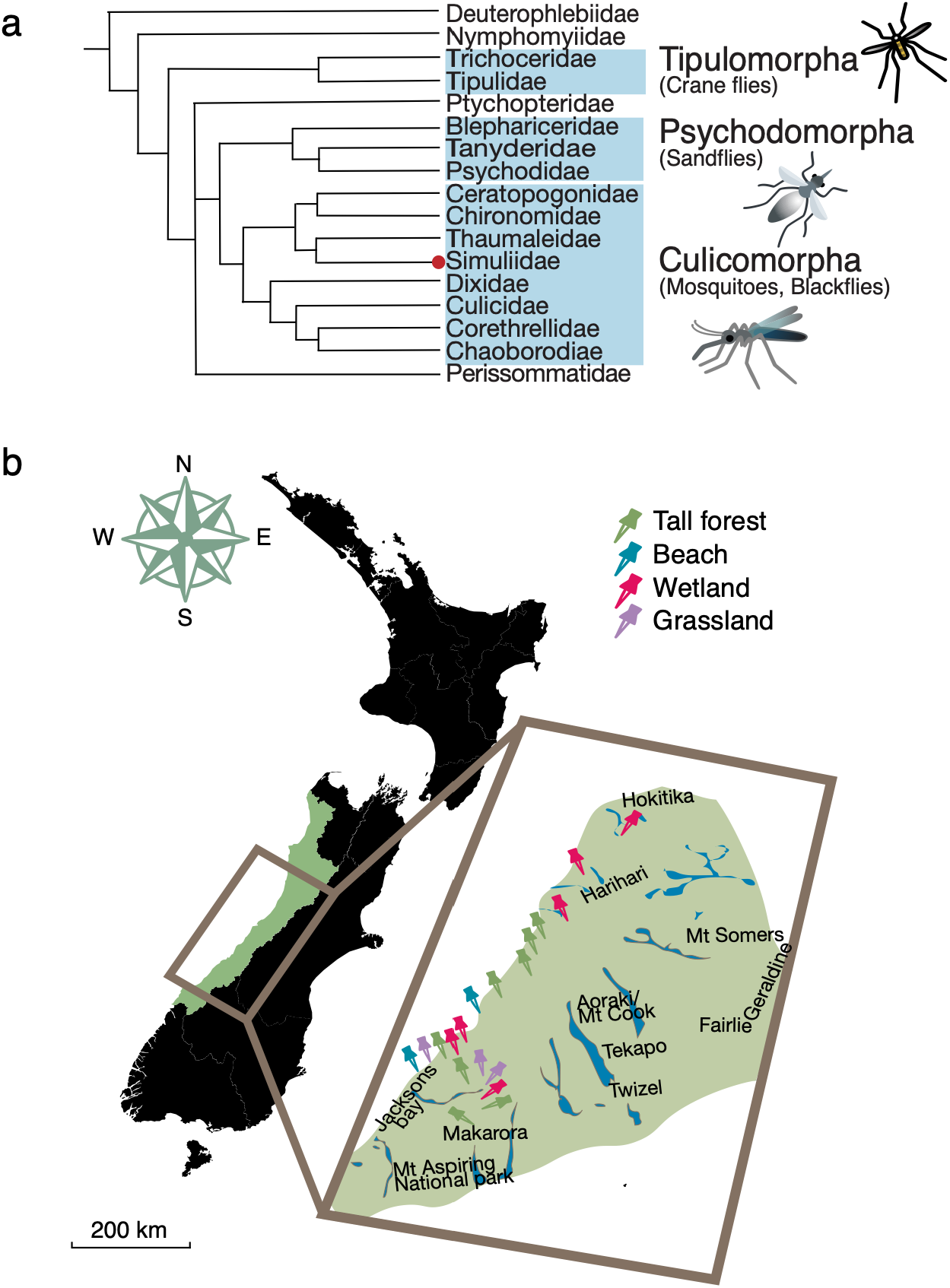
(**a**) Phylogeny of the families within infraorders Culicomorpha, Psychodomorpha and Tipulomorpha (42). The family Psychodidae contains phlebotomine sandflies which are shown to be phylogenetically distinct to New Zealand sandflies within the Simuliidae blackfly family (red dot). (**b**) Map of New Zealand with enlarged area of West Coast region. Each pin indicates a location where samples were collected. Each sampling location was classified and coloured by land use type as indicated in the top right corner key.

Based on their geographical distribution and morphological appearance, specimens were provisionally identified as the West Coast blackfly, although species identity was not confirmed using molecular methods (26,27). Collected specimens were transferred into sterile 2mL tubes containing 1mL of DNA/RNA Shield (Zymo Research, USA). Samples were maintained on ice packs during transport and transferred to −80 °C storage at the University of Otago, Dunedin, within three days of collection, where they remained until RNA extraction.

### RNA extraction and sequencing

Total RNA was extracted using the RNeasy Plus Mini Kit (QIAGEN, Germany), following the manufacturer’s protocol. Briefly, three to five blackflies were pooled per extraction and homogenised using a TissueRuptor (QIAGEN) in lysis buffer supplemented with β-mercaptoethanol. Homogenates were centrifuged to remove cellular debris, and the resulting supernatants were passed through gDNA eliminator spin columns to remove genomic DNA. RNA was subsequently purified using RNeasy spin columns, washed according to the manufacturer’s instructions and eluted in 40 µL of RNase-free water.

RNA concentration and purity were assessed using a NanoDrop spectrophotometer (Thermo Fisher Scientific, USA). Extracted RNA was combined into 20 sequencing pools representing 19 geographical locations, with two independent pools originating from Copland Valley (S4COP and S5COP). Equal volumes of individual RNA extracts were combined within each pool, resulting in final pooled volumes of 30–90 µL (Supplementary Table 1). Pooled RNA was submitted to the Australian Genome Research Facility (AGRF), Melbourne, Australia, for library preparation and sequencing. Following ribosomal RNA depletion, libraries were sequenced on an Illumina NovaSeq X platform to generate 150-bp paired-end reads.

### Metatranscriptomic assembly and virus identification

Raw sequencing reads were assessed for quality using FastQC (30), followed by adapter removal and quality trimming using Trimmomatic (31). Quality-filtered reads were assembled de novo using MEGAHIT v1.2.9 (32). Assembled contigs were screened for viral sequences by comparing predicted protein-coding regions against the National Center for Biotechnology Information (NCBI) protein database using DIAMOND BLAST v2.1.9 (33), with an E-value threshold of 1 × 10⁻¹⁰.

Putative viral contigs were further examined in Geneious Prime v2026.1 to identify and translate open reading frames (ORFs). Viral taxonomic assignments were informed by sequence similarity to characterised viruses, conserved protein domains and phylogenetic relationships. Contigs with matches to non-viral sequences or viruses unlikely to be associated with eukaryotic hosts were excluded from subsequent analyses.

To estimate viral abundance, quality-filtered reads were mapped to assembled viral contigs using Bowtie2 v2.4.4 (34). Viral read counts were normalised to reads per million (RPM) to account for differences in sequencing depth among libraries. Taxonomic assignments, sequence similarity to the closest characterised relatives and normalised read abundances were compiled for downstream phylogenetic and ecological analyses.

### Phylogenetic analysis

The evolutionary relationships of identified viruses were inferred using conserved polymerase protein sequences, including RNA-dependent RNA polymerase (RdRp) for RNA viruses and DNA polymerase for DNA viruses, where applicable. Reference sequences representing the closest matches identified by BLASTp with additional representative viruses from relevant taxonomic groups, were retrieved from NCBI.

Amino acid sequences were aligned using MAFFT v7.490 (35) with the L-INS-i algorithm, and poorly aligned regions were removed using trimAl v1.5.0 with a gap threshold of 0.9 (36). Maximum-likelihood phylogenetic trees were inferred using IQ-TREE v2.4.0 (37) under the LG amino acid substitution model, with branch support assessed using 1,000 ultrafast bootstrap replicates (38). Phylogenetic trees were visualised and annotated using FigTree v1.4.4 (39) and Adobe Illustrator v30.5.

Phylogenetic placement was used to assess the likely ecological associations of identified viruses. Viruses clustering with previously characterised invertebrate-associated viruses were considered putatively associated with blackflies, whereas those clustering predominantly with vertebrate-, bacterial-, fungal- or environmentally associated viruses were excluded from subsequent blackfly virome analyses. These classifications were treated as provisional, as phylogenetic placement alone cannot establish the host of replication or distinguish active infection from viruses acquired through feeding or environmental exposure.

### Viral abundance and community composition

Statistical analyses and data visualisation were performed in R using RStudio v2026.05.0, with the tidyverse and ggplot2 packages (40). Viral abundance was expressed as reads per million (RPM), calculated as the number of reads mapping to a viral sequence divided by the total number of sequencing reads in the corresponding library, multiplied by 10⁶.

Viral community composition was characterised at the family and phylum levels. To account for differences in overall viral abundance among libraries, relative abundances were calculated by dividing the RPM of each viral taxon by the sum of viral RPM values within the corresponding library. Relative abundance profiles were visualised using heatmaps with log₁₀ transformation to improve the representation of low-abundance taxa.

Viral richness was assessed as the number of distinct viral taxa detected within each library, and differences among land-use categories were examined. Mean richness and standard deviations were calculated for each land-use category, with individual library values displayed to illustrate variation among samples.

### Viral diversity and ecological analyses

Alpha diversity was quantified using the Shannon diversity index, calculated from family-level viral abundance data using the vegan package in R (41). Differences in viral diversity among land-use categories were assessed using Kruskal–Wallis tests. Additional comparisons were conducted at the viral phylum level.

Variation in viral community composition among libraries (beta diversity) was quantified using Bray–Curtis dissimilarity calculated from normalised viral abundance data. Patterns of community similarity were visualised using non-metric multidimensional scaling (NMDS).

Differences in viral community composition among land-use categories were assessed using permutational multivariate analysis of variance (PERMANOVA), conducted separately at the family and phylum levels.

To investigate geographical structuring of viral communities, pairwise geographical distances between sampling locations were estimated in kilometres using Google Earth. These distances were compared with pairwise Bray-Curtis dissimilarities to assess whether viral community composition became increasingly dissimilar with geographical separation. The strength of this relationship was evaluated using a Mantel test.

### Viral nomenclature and assessment of novelty

Putative novel viruses were identified based on amino acid sequence divergence from their closest characterised relatives. Viruses exhibiting less than 90% amino acid identity across conserved polymerase proteins, including RdRp for RNA viruses and DNA polymerase for DNA viruses, were provisionally classified as novel. Proposed virus names were assigned according to sampling location and their closest taxonomic affiliations. These designations are provisional and do not represent formal taxonomic assignments by the International Committee on Taxonomy of Viruses.

## Results

Metatranscriptomic sequencing of 20 pooled blackfly libraries generated between 84.5 million (S7FXG) and 161 million (S14MPL) paired-end reads per library (Figure 2a). Viral phylum richness did not differ significantly among the four land-use categories (p > 0.05 for all pairwise comparisons; Figure 2b), suggesting that the broad taxonomic diversity of viruses recovered was comparable across the sampled environments.

**Figure 2.**
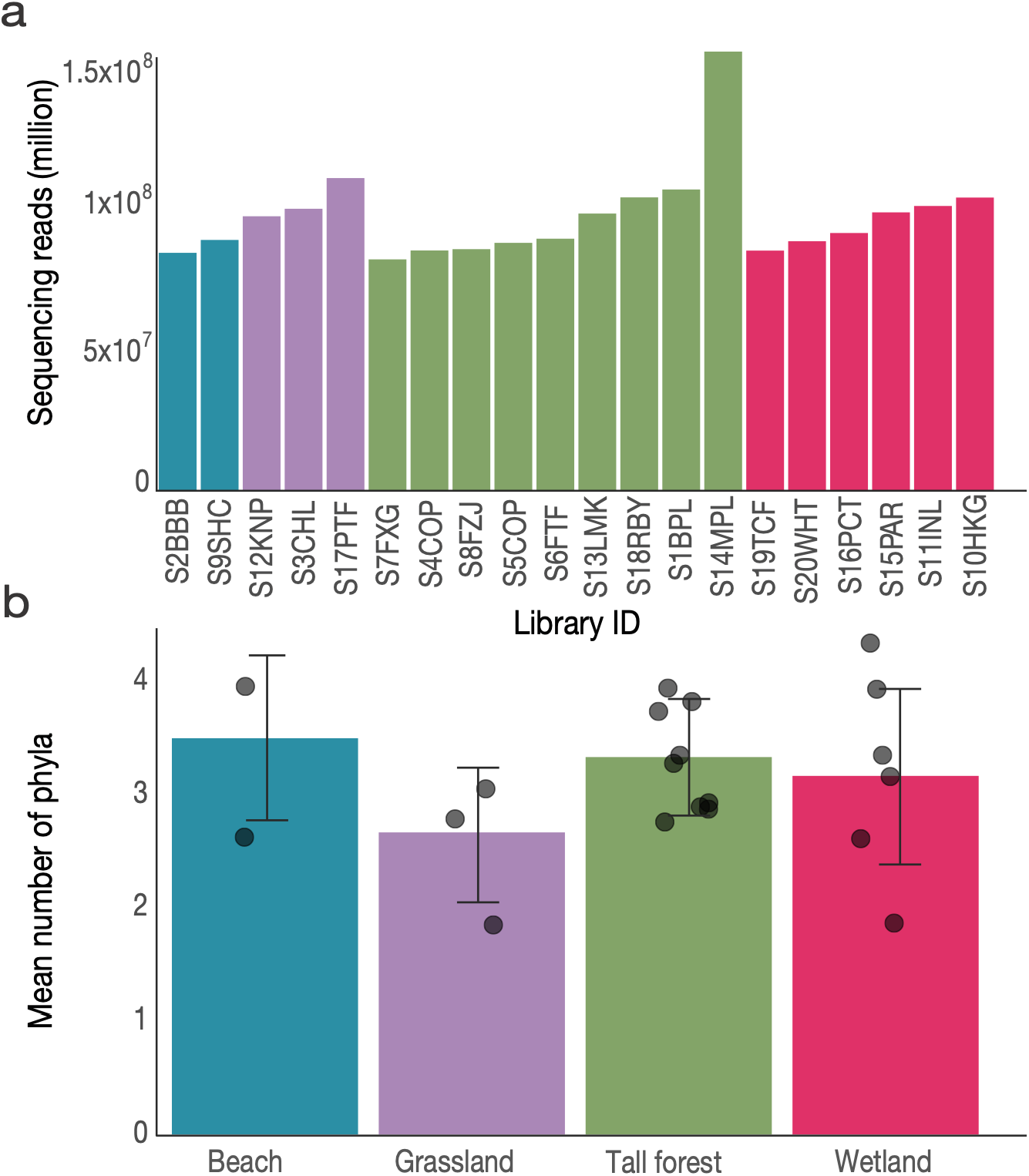
(**a**) Total paired-end sequencing reads generated for each library, coloured according to the land-use category of the sampling location. (**b**) Mean viral phylum richness across land-use categories. Individual points represent single libraries, and error bars indicate standard deviation.

Across all libraries, we identified 94 viruses spanning 15 viral families and six viral phyla, with two additional species classified only to the order level (Supplementary Table 2). All of the viruses identified were classified as putatively novel based on sequence divergence from their closest characterised relatives. The majority of viruses phylogenetically clustered with viruses previously identified from arthropods, particularly dipteran insects, although several clustered with viruses recovered from other hosts or environmental metagenomic studies.

The relative abundance of viral families varied considerably among libraries, with *Rhabdoviridae* and *Solemoviridae* collectively accounting for the greatest proportion of viral reads across the dataset (Figure 3). Although both families were detected across multiple land-use categories, their relative contributions differed markedly among environments. *Rhabdoviridae* accounted for 75.4% of viral relative abundance in beach habitats but only 7.5% in tall forest habitats. Conversely, *Solemoviridae* represented 83.8% of viral relative abundance in grassland habitats compared with 26.8% in wetlands. However, only two libraries originated from beach habitats, limiting the interpretation of differences involving this category.

**Figure 3.**
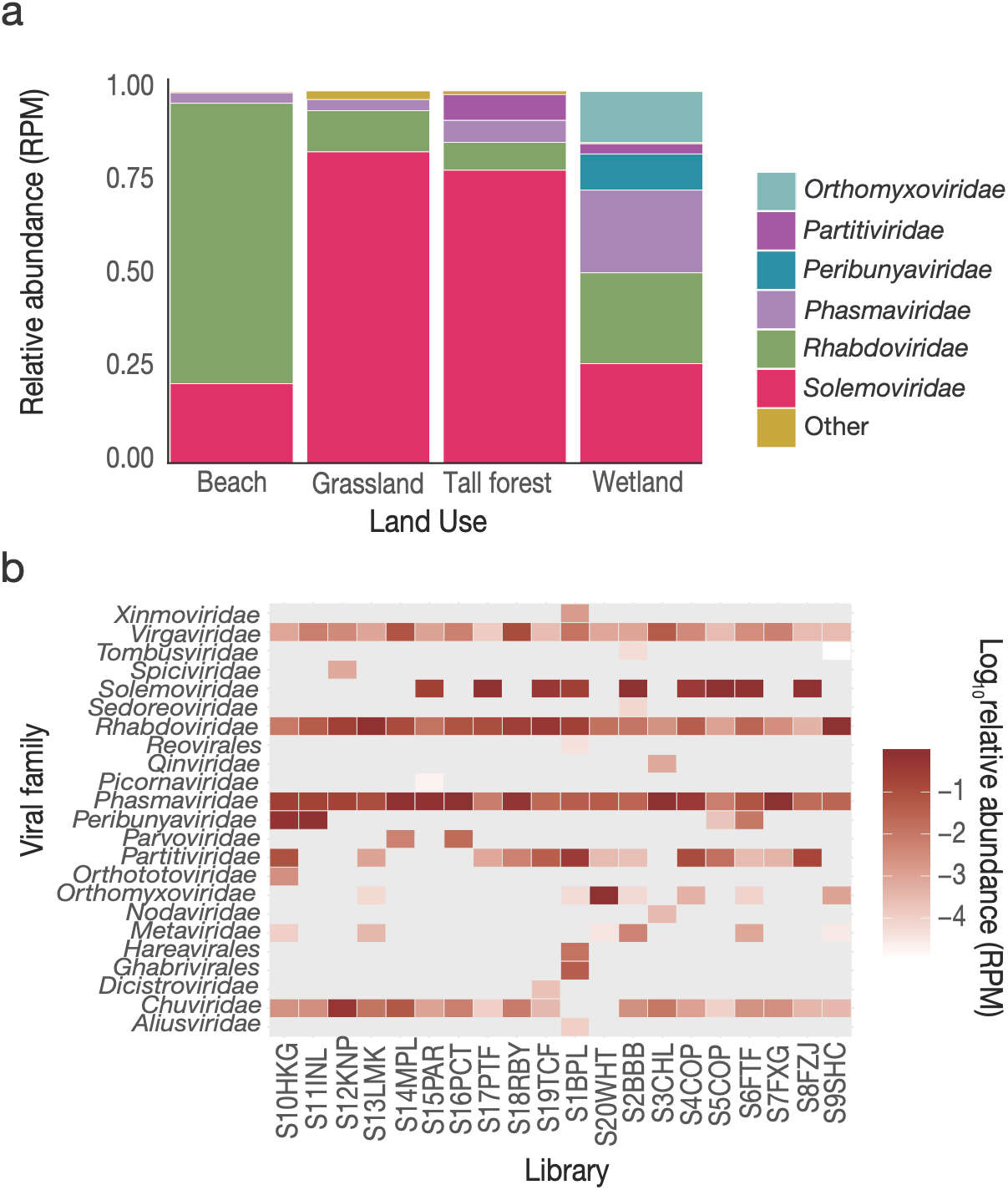
(**a**) Stacked bar plot showing the relative abundance of viral families across libraries grouped by land-use category. Relative abundance was calculated as the proportion of total viral RPM assigned to each family. Viral families contributing less than 2% of the total abundance were grouped as "Other". (**b**) Heatmap showing the abundance of viral families across individual libraries. Darker colours indicate higher abundance as indicated in the colour bar.

Several libraries were dominated by a single viral family, whereas others exhibited more even distributions of viral taxa, indicating substantial heterogeneity in viral community composition among sampling locations (Figure 3a). Members of *Rhabdoviridae* and *Phasmaviridae* were detected across numerous libraries, frequently at relatively high abundance. In contrast, several other viral families occurred at low relative abundance or were restricted to a small number of libraries (Figure 3b). These patterns indicate that the West Coast blackfly virome comprises both widely detected viral groups and taxa with more restricted distributions across the sampled locations.

To investigate the ecological structuring of blackfly viromes, we compared viral alpha and beta diversity among four land-use categories (Figure 4). Shannon diversity did not differ significantly among land-use types at either the viral family level (Kruskal–Wallis, χ² = 5.15, p = 0.161; Fig. 4a) or phylum level (χ² = 1.41, p = 0.704; Figure 4b). Thus, despite substantial differences in the relative abundance of individual viral families, there was no consistent association between land use and overall viral diversity.

**Figure 4.**
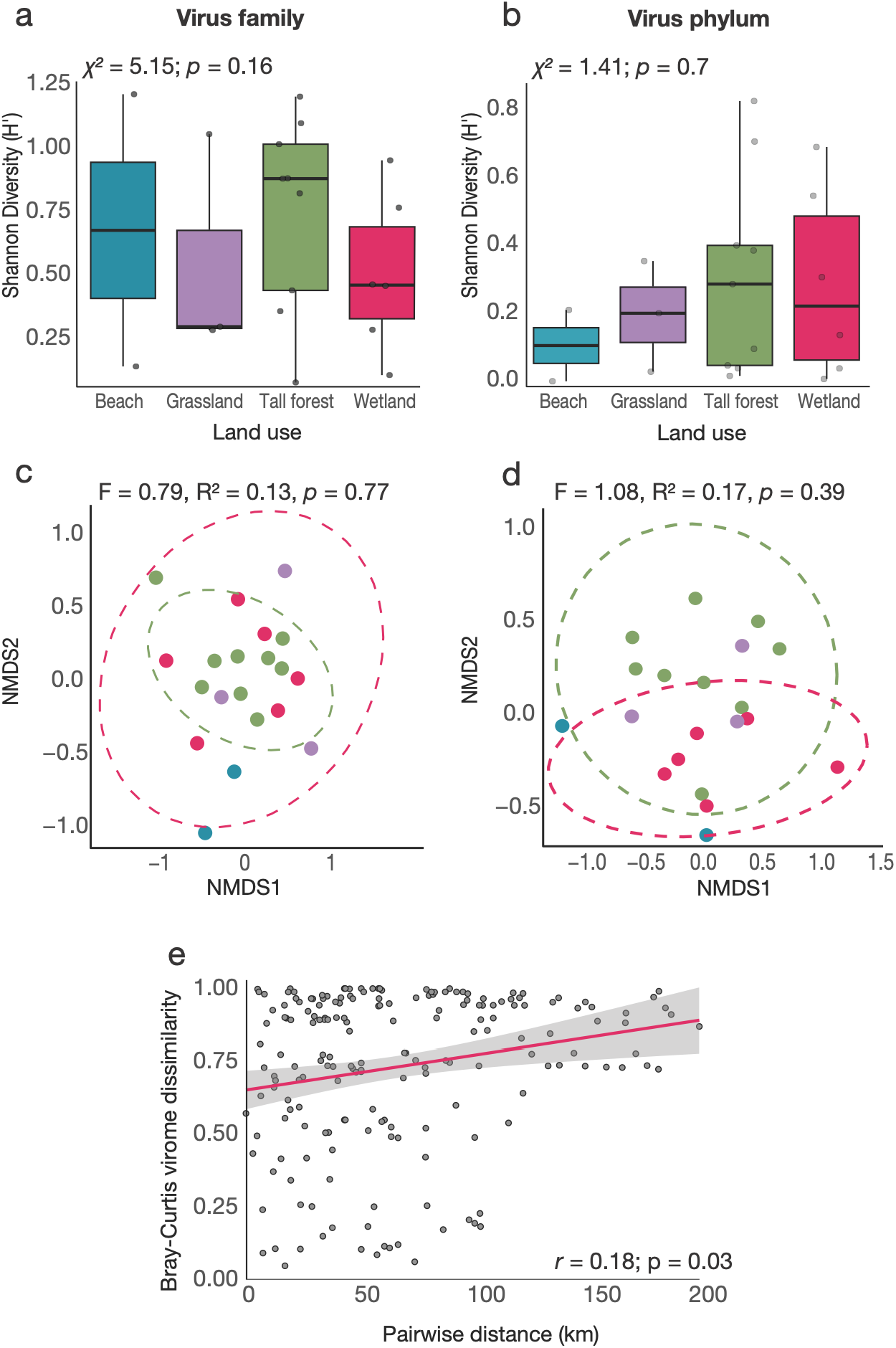
(**a**) Shannon diversity at the viral family level. (**b**) Shannon diversity at the viral phylum level. Boxplots show the median and interquartile range, with individual libraries shown as points. (**c**) NMDS ordination of Bray-Curtis dissimilarities at the viral family level. (**d**) NMDS ordination at the viral phylum level. Each point represents an individual library, with distances between points reflecting differences in viral community composition. (**e**) Relationship between pairwise geographic distance and Bray-Curtis dissimilarity of viral communities. Each point represents a pairwise comparison between sampling sites. A Mantel test identified a weak positive correlation between geographical distance and viral community dissimilarity (r = 0.18, p = 0.03).

Similarly, viral community composition did not differ significantly among land-use categories. At the family level, permutational multivariate analysis of variance (PERMANOVA) detected no significant association between land use and Bray-Curtis dissimilarity (F = 0.794, R² = 0.130, p = 0.774), with land use accounting for approximately 13% of the observed variation in community composition. Comparable results were obtained at the phylum level (F = 1.08, R² = 0.169, p = 0.394), where land use accounted for approximately 17% of the variation.

NMDS revealed substantial overlap among libraries from different land-use categories at both taxonomic levels (Figure 4c,d). Rather than forming distinct habitat-associated clusters, viral communities exhibited considerable variation among individual libraries, irrespective of land use. These findings suggest that the land-use categories examined were not strongly associated with viral community structure across the sampled locations, although the limited representation of some habitats, particularly beaches (n = 2 libraries), reduced the power to detect potential differences.

To determine whether geographical separation was associated with differences in viral community composition, we compared pairwise geographical distances between sampling locations with Bray-Curtis dissimilarities (Figure 4e). A positive correlation was observed between geographical distance and viral community dissimilarity (Mantel’s r = 0.18, p = 0.03).

Phylogenetic analyses of conserved viral polymerase sequences revealed substantial evolutionary diversity among the viruses recovered from West Coast blackflies. The 94 identified viruses were distributed across six viral phyla and 32 species, all of which were novel (Supplementary Table 2). Many occupied divergent phylogenetic positions relative to characterised viruses, including lineages associated with arthropods and environmental metagenomic datasets.

A single virus within *Cossaviricota*, provisionally named West Coast blackfly parvovirus, clustered within *Parvoviridae* but was distinct from known representatives, suggesting a putatively novel parvovirus (Figure 5). The greatest diversity occurred within *Negarnaviricota*, comprising six families (*Rhabdoviridae*, *Xinmoviridae*, *Chuviridae*, *Quinviridae*, *Orthomyxoviridae* and *Phasmaviridae*) and an unclassified lineage within *Hareavirales*. Rhabdoviruses were among the most abundant and widely distributed viruses, occurring across multiple libraries and clustering throughout the diversity of known arthropod-associated rhabdoviruses rather than forming a single monophyletic group. Several highly divergent xinmoviruses were also recovered across multiple libraries, clustering most closely with viruses previously identified from dipteran insects. Chuviruses and quinviruses formed distinct lineages within their respective families, although quinviruses were generally less abundant. A single orthomyxovirus clustered with arthropod-associated rather than vertebrate-associated viruses, while an additional divergent virus within *Hareavirales* could not be assigned to a recognised family. Phasmaviruses were also abundant and detected across geographically separated libraries, forming distinct lineages within the known diversity of insect-associated phasmaviruses.

**Figure 5.**
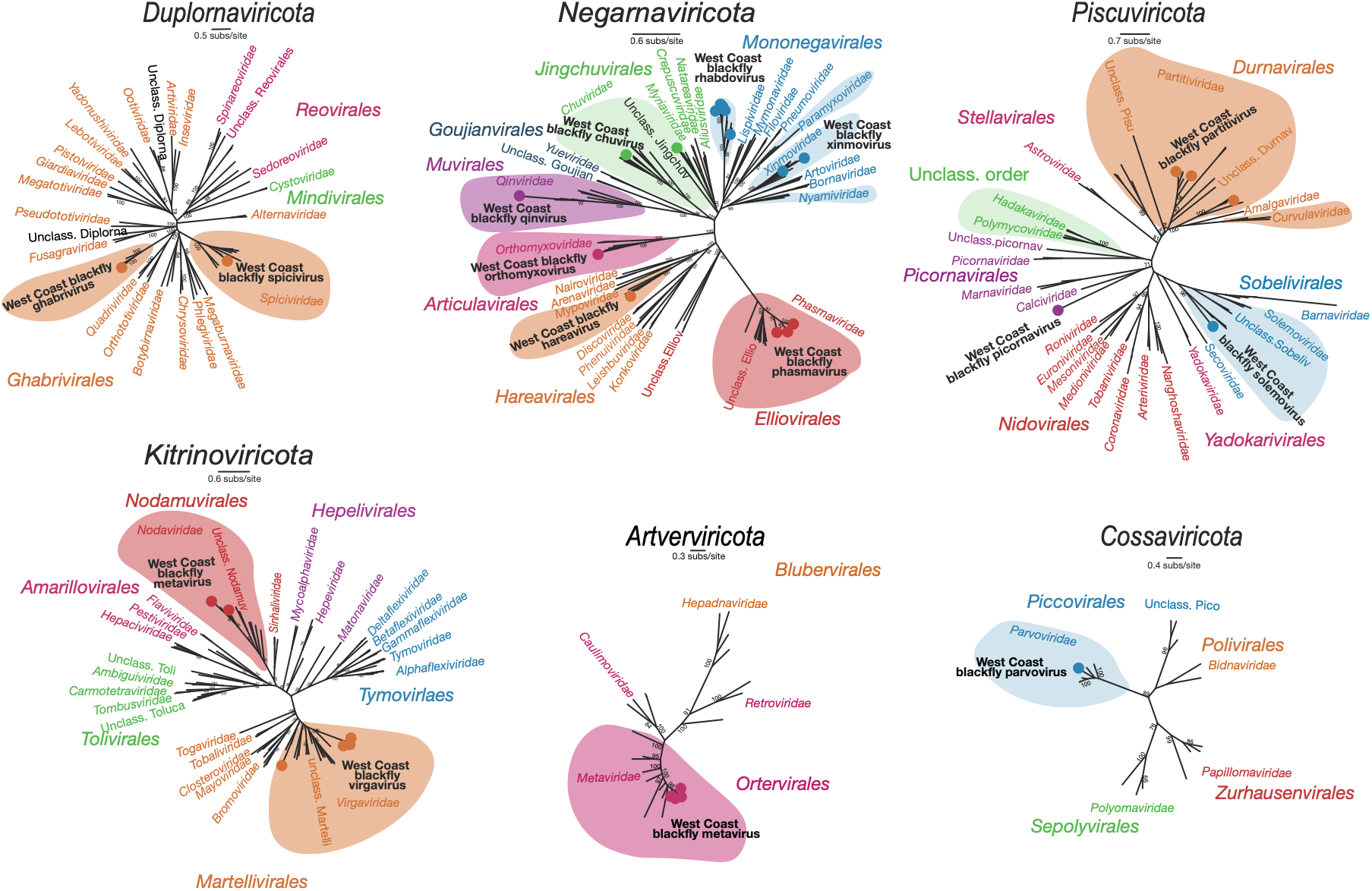
Maximum likelihood phylogenetic trees of viruses representing six viral phyla identified in West Coast blackflies. Unrooted maximum likelihood phylogenetic trees of representative viruses from the phyla: *Cossaviricota, Negarnaviricota, Kitrinoviricota, Pisuviricota, Artverviricota* and *Duplornaviricota.* Phylogenetic relationships were inferred from conserved amino acid sequences of DNA polymerase (*Cossaviricota*), RNA-dependent RNA polymerase (*Negarnaviricota*, *Kitrinoviricota*, *Pisuviricota* and *Duplornaviricota*), or reverse transcriptase (*Artverviricota*). Viruses identified in this study are highlighted in bold and with a coloured circle at the tip. Bootstrap support values greater than 70% are shown at major nodes. The families *Belpaoviridae* and *Pseudoviridae* were excluded from the *Artverviricota* phylogeny because representative conserved gene sequences were unavailable.

Viruses within *Kitrinoviricota* belonged to *Virgaviridae* and *Nodaviridae* (Figure 5). Multiple closely related but divergent virgaviruses, provisionally named West Coast blackfly virgaviruses, formed well-supported lineages distinct from characterised representatives. Two nodaviruses clustered with known insect-associated nodaviruses but occupied a distinct phylogenetic position, consistent with a putatively novel species.

The phylum *Pisuviricota* comprised viruses from *Solemoviridae*, *Partitiviridae* and *Picornaviridae* (Figure 5). Solemoviruses were among the most abundant viruses recovered and were detected in most libraries, with several forming divergent lineages relative to characterised species. Partitiviruses were less abundant but occurred across multiple libraries and were phylogenetically distinct from known representatives. Given their established associations with plants, fungi and protists, their detection may reflect dietary or environmental sources rather than infection of blackflies. Picornaviruses were detected less frequently and at relatively low abundance, forming distinct branches within the recognised diversity of the family.

Several viruses within *Artverviricota* were assigned to *Metaviridae* and provisionally named West Coast blackfly metaviruses. These were recovered across multiple libraries, with phylogenetic analysis of reverse transcriptase sequences revealing several divergent lineages within the family. Other recognised families, including *Belpaoviridae* and *Pseudoviridae*, were excluded from the phylogenetic reconstruction because suitable reference sequences were unavailable.

Finally, *Duplornaviricota* was represented by two divergent viruses detected at relatively low abundance (Figure 5). One clustered within *Spiciviridae*, forming a distinct lineage consistent with a putatively novel double-stranded RNA virus, while the other was placed within *Ghabrivirales* but could not be confidently assigned to an existing family.

## Discussion

Metatranscriptomic sequencing of West Coast blackflies from Aotearoa New Zealand revealed extensive viral diversity, including 94 viruses across 32 species, all of which were putatively novel. These findings add to growing evidence that arthropods harbour a substantial proportion of the largely unexplored RNA virosphere, with many viral lineages remaining poorly represented in existing sequence databases (7, 8, 44–46). The recovery of highly divergent viruses across multiple phyla, including lineages with distant relationships to previously characterised arthropod-associated viruses, highlights the evolutionary diversity associated with a single endemic insect lineage. These findings demonstrate how virome investigations of geographically isolated and historically understudied arthropods can expand our understanding of viral evolution and host associations beyond the relatively small number of well-characterised vector species.

The greatest viral diversity was observed within *Negarnaviricota*, particularly among members of *Rhabdoviridae*, *Xinmoviridae*, *Chuviridae* and *Phasmaviridae*. Rhabdoviruses were consistently detected at high abundance across multiple sampling locations, including three putatively novel viruses (Supplementary Table 2). Rather than forming a single monophyletic group, these viruses were distributed across the broader phylogenetic diversity of arthropod-associated rhabdoviruses, suggesting that blackflies are associated with multiple evolutionarily distinct viral lineages. The identification of three putatively novel phasmaviruses across geographically separated locations similarly suggests that members of this predominantly insect-associated family are widespread within the sampled blackfly populations. Highly divergent viruses within *Xinmoviridae* and *Chuviridae* further extend the known diversity of these relatively poorly characterised groups (44, 47).

Several viruses were also phylogenetically associated with families containing recognised vertebrate pathogens, including *Rhabdoviridae* and *Orthomyxoviridae*. Rhabdoviruses encompass a broad range of vertebrate- and invertebrate-associated viruses, including vesicular stomatitis virus, for which blackflies have demonstrated vector competence under experimental conditions (21, 48). Similarly, some arthropod-associated orthomyxoviruses, including quaranjaviruses, have been associated with birds and ticks (49, 50). However, membership of these viral families does not imply vertebrate infectivity or transmission potential. Indeed, the orthomyxovirus identified here clustered with arthropod-associated rather than vertebrate-associated lineages. Moreover, viruses detected in whole insects may originate from blood meals, associated microorganisms or environmental material rather than active infection of blackflies (8, 10, 55). Establishing host associations and transmission potential would therefore require additional evidence of viral replication, tissue tropism and, where relevant, vector competence. These distinctions are particularly important when interpreting highly divergent viruses whose biological properties cannot be reliably inferred from phylogenetic relationships alone.

Beyond virus discovery, our findings provide insights into the spatial organisation of arthropod-associated viral communities. Viral community composition varied substantially among sampling locations, with a positive relationship between geographical distance and virome dissimilarity across an approximately 187 km transect. Geographical and environmental structuring of viral communities has been described across diverse ecosystems, including marine environments and arthropod-associated viromes (51, 52). Such patterns may arise through spatially restricted host movement, differences in local environmental conditions or variation in the availability of vertebrate hosts and other ecological resources. The relatively restricted geographical extent of our sampling makes this observation particularly interesting, suggesting that spatial structuring of viral communities may occur even across comparatively small landscapes.

In contrast, broad land-use categories were not significantly associated with viral richness, Shannon diversity or community composition. Although the relative abundance of individual viral families differed markedly among habitats, these differences did not translate into consistent community-level patterns. Previous studies have demonstrated that land use, vegetation, climate and host ecology can influence the composition of arthropod-associated viromes (43, 52, 53). However, the absence of a detectable association in this study suggests that broad habitat classifications may not adequately capture the ecological factors shaping viral communities in West Coast blackflies. Local environmental conditions, host availability, insect population structure and stochastic variation in viral abundance may be more informative.

Several additional factors should be considered when interpreting these findings. Sampling was conducted over a restricted period, limiting our ability to assess seasonal or temporal variation in viral diversity. Viral identification and taxonomic assignment remain constrained by the uneven representation of host species and geographical regions in public sequence databases (55, 56). Highly divergent viral sequences may lack sufficiently close relatives for confident classification, contributing to the substantial unexplored sequence diversity commonly described as viral ’dark matter’ (54–56). Consequently, the designation of putatively novel viruses and their inferred host associations should be considered provisional, particularly where only partial genomic sequences were recovered.

In conclusion, our findings reveal substantial and previously undescribed viral diversity associated with an endemic blackfly lineage and suggest that viral community composition may exhibit spatial structuring even across relatively restricted geographical scales. By extending virus discovery into an understudied arthropod group, this study contributes to a broader understanding of viral diversity and the ecological processes governing its distribution.

## Supplementary information

**Supplementary Table 1. Metadata of RNA pools.** Table of 20 pools based on location and associated metadata. The location of sample collection for each pool is noted with exact coordinates of sampling site. Land use of each site is based on the New Zealand Ministry for the Environment LUCAS land use map (available at https://environment.govt.nz/facts-and-science/science-and-data/new-zealand-land-use-map/, accessed November 2026).

**Supplementary Table 2. Novel viral transcripts identified in this study.** Taxonomic classification and provisional names assigned to each novel viral transcript, together with its closest known relative and the number of sequencing libraries in which it was detected.

## Data availability

All virus sequences identified are available on NCBI’s Genbank database under accession numbers QB082497-QB082587 and QB146866-QB146868. The raw reads generated are available on the Sequence Read Archive (SRA) under BioProject PRJNA1538114.

## Author contributions

Jemma Geoghegan designed the study. Kimiya Byrne and Jessica Darnley conducted the fieldwork and sample collection. Kimiya Byrne performed the laboratory work, led bioinformatic analyses and prepared the first manuscript draft. Stephanie Waller supervised the laboratory and bioinformatic analyses. Jemma Geoghegan secured funding and overall supervision. All authors contributed to reviewing and editing the manuscript and approved the final version.

