## Supplementary Table 1 for "Novel viral diversity and geographic structuring in the New Zealand West Coast blackfly (*Austrosimulium ungulatum*)"

**Supplementary Table 1.** **Metadata of RNA pools.** Table of 20 pools based on location and associated metadata. The location of sample collection for each pool is noted with exact coordinates of sampling site. Land use of each site is based on the New Zealand Ministry for the Environment LUCAS land use map (available at <https://environment.govt.nz/facts-and-science/science-and-data/new-zealand-land-use-map/>, accessed November 2026).

| **Pool ID** | **Location** | **Latitude** | **Longitude** | **Land use** | **#sandflies collected** | **Date of Collection** |
| --- | --- | --- | --- | --- | --- | --- |
| S1BPL | Blue Pools | 44°09’34.0"S | 169°17'05.9"E | Tall forest | 22 | 13/11/25 |
| S2BBB | Bruce Bay | 43°35’41.0"S | 169°36'15.5"E | Beach | 27 | 13/11/25 |
| S3CHL | Confluence of Haast and Landsborough rivers | 43°58’39.1"S | 169°25'19.2"E | Grassland | 23 | 13/11/25 |
| S4COP | Copland Valley | 43°34’28.9"S | 169°48'41.6"E | Tall forest | 38 | 13/11/25 |
| S5COP | Copland Valley | 43°34’28.9"S | 169°48'41.6"E | Tall forest | 40 | 13/11/25 |
| S6FTF | Fantail Falls | 44°04’40.3"S | 169°23'08.2"E | Tall forest | 19 | 13/11/25 |
| S7FXG | Fox Glacier | 43°28’45.6"S | 170°00'30.8"E | Tall forest | 4 | 13/11/25 |
| S8FZJ | Franz Josef | 43°22’21.5"S | 170°10'47.6"E | Tall forest | 14 | 13/11/25 |
| S9SHC | Ships Creek Haast | 43°45’28.2"S | 169°08'52.4"E | Beach | 12 | 10/11/25 |
| S10HKG | Hokitika Gorge | 42°57'18.2"S | 171°00'58.3"E | Wetland | 20 | 12/11/25 |
| S11INL | Ianthe Lake | 43°03’35.2"S | 170°38'02.5"E | Wetland | 12 | 12/11/25 |
| S12KNP | Knights  Point | 43°42’54.2"S | 169°13'32.3"E | Grassland | 22 | 13/11/25 |
| S13LMK | Lake Moeraki | 43°43’56.6"S | 169°18'03.9"E | Tall forest | 22 | 13/11/25 |
| S14MPL | Lake Mapourika | 43°19’44.5"S | 170°12'50.1"E | Tall forest | 23 | 12/11/25 |
| S15PAR | Paringa Campsite | 43°43’18.1"S | 169°24'39.3"E | Wetland | 25 | 13/11/25 |
| S16PCT | Paringa Cattle Track | 43°46’20.1"S | 169°21'17.6"E | Wetland | 23 | 13/11/25 |
| S17PTF | Pleasant Flat | 44°00’43.2"S | 169°22'52.6"E | Grassland | 26 | 13/11/25 |
| S18RBY | Roaring Billy Falls | 43°56’19.4"S | 169°17'07.2"E | Tall forest | 21 | 13/11/25 |
| S19TCF | Thunder Creek Falls | 44°02’13.5"S | 169°22'01.9"E | Wetland | 22 | 13/11/25 |
| S20WHT | Whataroa | 43°17’37.2"S | 170°24'44.9"E | Wetland | 10 | 12/11/25 |
