## Supplementary Table 2 for "Novel viral diversity and geographic structuring in the New Zealand West Coast blackfly (*Austrosimulium ungulatum*)"

**Supplementary Table 2. Novel viral transcripts identified in this study.** Taxonomic classification and provisional names assigned to each novel viral transcript, together with its closest known relative and the number of sequencing libraries in which it was detected.

| *Virus (proposed name)* | *Phylum* | *Family* | *Closest relative in NCBI GenBank* | *# Libraries detected* |
| --- | --- | --- | --- | --- |
| West Coast blackfly parvovirus | Cossaviricota | *Parvoviridae* | Parvoviridae sp. | 1 |
| West Coast blackfly rhabdovirus 1 | Negarnaviricota | *Rhabdoviridae* | Leksozero Simulium Almendravirus | 5 |
| West Coast blackfly rhabdovirus 2 | Negarnaviricota | *Rhabdoviridae* | Leksozero Simulium Almendravirus | 1 |
| West Coast blackfly rhabdovirus 3 | Negarnaviricota | *Rhabdoviridae* | Riboviria sp. | 1 |
| West Coast blackfly xinmovirus 1 | Negarnaviricota | *Xinmoviridae* | Spider associated mononega virus 10 | 1 |
| West Coast blackfly xinmovirus 2 | Negarnaviricota | *Xinmoviridae* | Aedes albopictus anphevirus1 | 1 |
| West Coast blackfly chuvirus 1 | Negarnaviricota | *Chuviridae* | Ophiovirus sp. ishikawaense | 1 |
| West Coast blackfly chuvirus 2 | Negarnaviricota | *Chuviridae* | Shuangao Fly Virus 1 | 1 |
| West Coast blackfly qinvirus | Negarnaviricota | *Qinviridae* | Palkane qinvirus | 1 |
| West Coast blackfly orthomyxovirus | Negarnaviricota | *Orthomyxoviridae* | Dipteran orthomyxo-related virus OKIAV193 | 1 |
| West Coast blackfly hareavirus | Negarnaviricota | *Order: Hareavirales* | Dipteran mypo-related virus OKIAV322 | 1 |
| West Coast blackfly phasmavirus 1 | Negarnaviricota | *Phasmaviridae* | Dipteran phasma-related virus OKIAV225 | 19 |
| West Coast blackfly phasmavirus 2 | Negarnaviricota | *Phasmaviridae* | Dipteran phasma-related virus OKIAV225 | 20 |
| West Coast blackfly phasmavirus 3 | Negarnaviricota | *Phasmaviridae* | Barstukas virus | 1 |
| West Coast blackfly nodavirus 1 | Kitrinoviricota | *Nodaviridae* | Nika virus | 1 |
| West Coast blackfly nodavirus 2 | Kitrinoviricota | *Nodaviridae* | Army ant associated nodavirus 2 | 1 |
| West Coast blackfly virgavirus 1 | Kitrinoviricota | *Virgaviridae* | Xanka Hybomitra Negev-like virus | 12 |
| West Coast blackfly virgavirus 2 | Kitrinoviricota | *Virgaviridae* | Bemisia tabaci virga-like virus 1 | 2 |
| West Coast blackfly virgavirus 3 | Kitrinoviricota | *Virgaviridae* | Bemisia tabaci virga-like virus 1 | 1 |
| West Coast blackfly virgavirus 4 | Kitrinoviricota | *Virgaviridae* | Asiatic toad virga-like virus 3 | 1 |
| West Coast blackfly partitivirus 1 | Pisuviricota | *Partitiviridae* | Soybean thrips partiti-like virus 10 | 6 |
| West Coast blackfly partitivirus 2 | Pisuviricota | *Partitiviridae* | Riboviria sp. | 1 |
| West Coast blackfly partitivirus 3 | Pisuviricota | *Partitiviridae* | Spider partiti-like virus 285 | 1 |
| West Coast blackfly picornavirus | Pisuviricota | *Picornaviridae* | Guangdong ant virus 2 | 1 |
| West Coast blackfly solemovirus 1 | Pisuviricota | *Solemoviridae* | Riboviria sp. | 3 |
| West Coast blackfly solemovirus 2 | Pisuviricota | *Solemoviridae* | Virus sp. | 1 |
| West Coast blackfly metavirus 1 | Artverviricota | *Metaviridae* | Pitica errantivirus | 2 |
| West Coast blackfly metavirus 2 | Artverviricota | *Metaviridae* | *Aedes aegypti* To virus 1 | 2 |
| West Coast blackfly metavirus 3 | Artverviricota | *Metaviridae* | *Aedes aegypti* To virus 1 | 1 |
| West Coast blackfly metavirus 4 | Artverviricota | *Metaviridae* | *Anopheles coluzzii* | 1 |
| West Coast blackfly ghabrivirus | Duplornaviricota | *Order:*  *Ghabrivirales* | Sopuglig virus | 1 |
| West Coast blackfly spicivirus | Duplornaviricota | *Spiciviridae* | Riboviria sp. | 1 |
